# ChipSeg: an automatic tool to segment bacteria and mammalian cells cultured in microfluidic devices

**DOI:** 10.1101/2020.08.03.225045

**Authors:** Irene de Cesare, Criseida G. Zamora-Chimal, Lorena Postiglione, Mahmoud Khazim, Elisa Pedone, Barbara Shannon, Gianfranco Fiore, Giansimone Perrino, Sara Napolitano, Diego di Bernardo, Nigel Savery, Claire Grierson, Mario di Bernardo, Lucia Marucci

## Abstract

Extracting quantitative measurements from time-lapse images is necessary in external feedback control applications, where segmentation results are used to inform control algorithms. While such image segmentation applications have been previously reported, there is in the literature a lack of open-source and documented code for the community. We describe ChipSeg, a computational tool to segment bacterial and mammalian cells cultured in microfluidic devices and imaged by time-lapse microscopy. The method is based on thresholding and uses the same core functions for both cell types. It allows to segment individual cells in high cell-density microfluidic devices, to quantify fluorescence protein expression over a time-lapse experiment and to track individual cells. ChipSeg enables robust segmentation in external feedback control experiments and can be easily customised for other experimental settings and research aims.

## Introduction

Live-cell imaging by automated microscopy enables the collection of large-scale data, useful to study the link between cellular dynamics and emerging phenotypes.

In the context of synthetic biology, the combination of control engineering algorithms with live-cell imaging has been shown to successfully enable the automatic regulation of gene expression across cellular chassis ^1–8^. If employing microfluidics/microscopy platforms, the external feedback control action is implemented by measuring the relevant control output (e.g. fluorescent reporter expression in cells grown within microfluidic devices) throughout the time-lapse experiment by means of automatic segmentation. This measurement informs a control algorithm that computes the control input to minimize the control error (i.e. the difference between the control target and output). The control input is automatically provided to cells, for example by changing the culture media with motor-controlled pumps. Automatic segmentation needs to be performed in real-time and to be robust over the whole time-lapse experiment.

We present here ChipSeg, a thresholding-based algorithm to automatically segment both bacterial and mammalian cells cultured in microfluidic devices. The same core functions are implemented for both cell types, making the code flexible for other chassis and applications. The algorithm showed robust segmentation results in external feedback control experiments with microfluidics/microscopy platforms ^6–9^, and can be easily adapted for open-loop experiments, other cell types and experimental settings.

ChipSeg is implemented in MATLAB and it is publicly available. The source code and documentation can be found at https://github.com/LM-group/ChipSeg.

## Results

ChipSeg is broadly based on the Otsu thresholding method ^10^ and is implemented in Matlab, using both custom and built-in Image Processing Toolbox functions. When used within an external feedback control experiment, the segmentation is performed online: at each sampling time, the acquired image is imported by ChipSeg and the algorithm output is fed-back to the computer for control input calculation (Fig. 1A, B).

**Figure 1.**
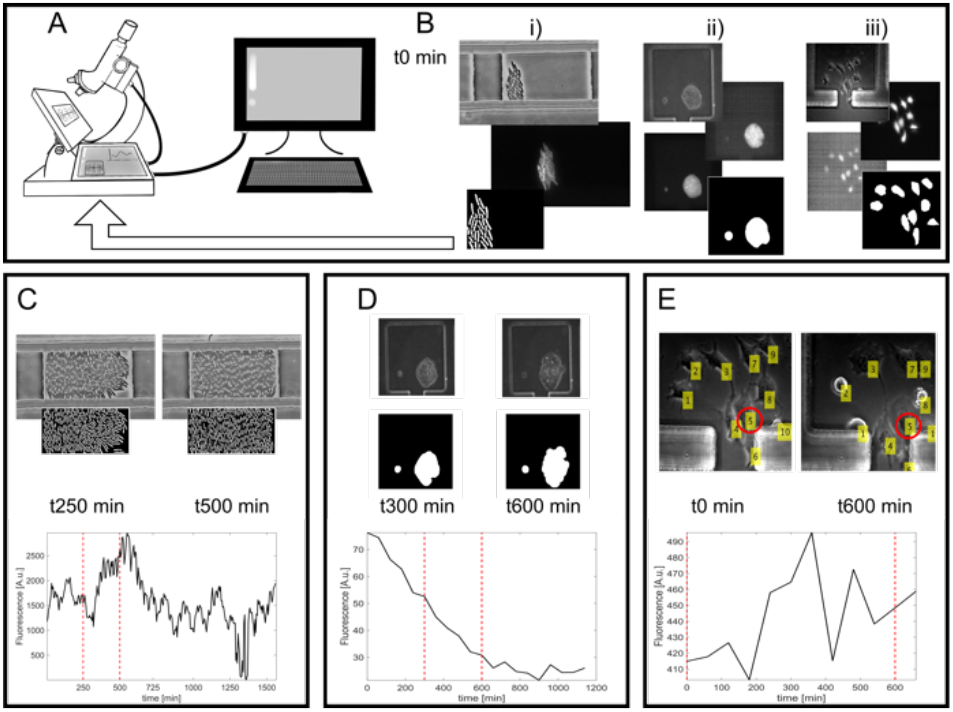
ChipSeg main features. **A, B)** The algorithm can segment time-lapse images of bacterial (B i) and mammalian cells with dome-shaped (B ii) or flat (B iii) morphology, cultured in microfluidic devices; if used within an external feedback control experiment, at each sampling time the image acquired by the microscope is segmented and ChipSeg output is fed-back to the computer, which can compute the control error and input. In B), phase-contrast, fluorescence channels -green in i), blue and green in ii), red and infrared in iii)- and ChipSeg-computed masks (i, ii) and ellipsoids (iii) are shown. Masks are computed on a cropped area of the acquired raw image. **C, D)** Average fluorescence quantification in exemplar bacterial cells (C) and Rex1-GFPd2 mouse embryonic stem cells (mESCs, D) time-lapses (Supporting Information); the dotted red lines indicate the time points for which the phase-contrast raw images and the corresponding ChipSeg-computed masks on a cropped region are shown. **E)** Two time points (dotted red lines) of a dual-reporter mESCs (Supporting Information) time-lapse, showing the overlay of phase-contrast images and the ChipSeg-computed single-cell label, tracked over time; quantification of the mCherry fluorescent reporter for a specific cell (labelled as 5) over the time-lapse is shown.

Firstly, ChipSeg performs global thresholding to segment a bacterial or mammalian cell cluster in a region of interest within the microfluidic device used for cell culturing; pre-processing, such as filtering and contrast enhancement, can be applied (Supporting Information). Global thresholding can be performed on phase-contrast images or, in case of poor contrast, on fluorescent channel images of cells stained with a fluorescent dye (Supporting Information).

If cells are growing as a monolayer, local thresholding is performed to segment individual cells (bacterial and mammalian cells in Fig. 1B i and iii, respectively); global thresholding only is performed if cells are growing in dome-shaped colonies (mammalian cells in Fig. 1B ii). The segmentation can be refined in a customized way to fix possible artefacts, for example by removing or separating objects smaller/bigger than a set threshold, respectively (Supporting Information).

Fluorescence is calculated by overlapping the segmented masks to the fluorescent channel image; in so doing, a binary mask defining the pixels of the fluorescence image is calculated. Fig. 1C, D and Supporting Movies 1 and 2 show the average fluorescence of cells within the selected region of interest for exemplar bacterial and mammalian cell time-lapses, respectively.

If local thresholding is performed, the algorithm provides the number of segmented cells, which can serve to estimate the cell population growth rate over the time-lapse experiment (Supporting Movie 1). Furthermore, by computing the maximum and minimum of fluorescence in the cellular population, cell-to-cell variability can be quantified.

ChipSeg also allows for cell tracking, provided movement across frames is limited and the microfluidic device is not too dense. The algorithm firstly computes a mask by applying filters and thresholding functions, then assigns a centroid to an individual cell, and finally searches the correspondence between individual cells in two consecutive time-lapse images by minimizing the centroid displacement (Fig. 1E, Supporting Movie 3).

Our algorithm is fully customizable, as we provide a step-by-step explanation of all the pre-processing, segmentation, tracking and fluorescence quantification functions (Supporting Information).

## Discussion

Many automated image segmentation tools exist to analyse microscopy images ^11–13^; user-friendly graphical interfaces within a software package can present difficulties at the time of interfacing them with other software (e.g., in external feedback control applications, where code which can tune the control action given the segmentation output is required).

Other image analysis algorithms previously proposed to segment cells cultured and imaged within microfluidic devices are often specific for certain chip designs such as the mother machine device ^14–16^; instead, ChipSeg does not rely on the geometry of the devices we used (see ^17^ and ^18^ for bacterial and mammalian cells device description, respectively).

ChipSeg is based on simple Matlab functions and should be easy to customise also for users lacking a computational background. While it is true that parameters within the algorithm might need to be tuned if using images of different cell lines or if varying acquisition settings, a few trial-and-error iterations might be enough to fix them, using limited images.

Our algorithm has enough flexibility to be used across cell types, and adaptation of the code should be quicker than implementing deep learning-based approaches (e.g. ^16, 19–21^), whose undeniable robustness, in most cases, comes at the cost of extensive algorithm training on manually annotated datasets.

## ASSOCIATED CONTENT

**Supporting Information.PDF** This file contains a step-by-step explanation of the functions used.

**Supporting Movie 1** Exemplar bacterial cell movie, showing phase-contrast and GFP fluorescent images, ChipSeg-computed mask, average fluorescence and cell number.

**Supporting Movie 2** Exemplar Rex1-GFPd2 mESC movie, showing phase-contrast and green/blue fluorescent images. ChipSeg-computed mask and fluorescence are reported.

**Supporting Movie 3** Exemplar dual-reporter mESC movie, showing phase-contrast and mCherry/infrared images. ChipSeg-computed cell labels and their tracking over time are shown, as well as mCherry fluorescence of an individual cell (labelled as 5) over the time-lapse.

## AUTHOR INFORMATION

### Corresponding Author

M.d.B. and L.M.

### Author Contributions

L.M. and M.d.B. conceived of the project and supervised the work; L.P., G.F., G.P. and S.N. developed the initial code; I.d.C. and C.G.Z.C. refined the algorithm and analysed time-lapses; M.K., B.S. and E.P. performed the experiments; D.d.B. supervised S.N. and G.P.; N.S. and C.G. co-supervised the experimental activities with bacteria; I.d.C., C.G.Z.C. and L.M. wrote the paper with input from the other authors.

## ACKNOWLEDGMENT

We thank Dr Mark Jepson and Alan Leard (Wolfson Imaging Facility, University of Bristol) for supporting live-cell imaging experiments.

