## Supporting information for "ChipSeg: an automatic tool to segment bacteria and mammalian cells cultured in microfluidic devices"

### Bacterial cell segmentation

The segmentation algorithm is applied here to phase contrast images to distinguish the foreground (bacteria cells) from the background in each frame of the time-lapse experiment. A pre-processing of the raw images is generally required before running the segmentation routine, which can then segment and count individual cells in a filtered frame. The segmentation algorithm output is a mask, used to calculate the fluorescence of individual cells and the average fluorescence across the bacteria population. The Matlab files implementing the reported algorithm are *'segmentation_main_offline_ol_video.m'*, *'segmentation_GF.m', 'fluorescence_eval_FG_Init.m', 'fluorescence_eval_GF.m' and reshapeHist.m.*

#### 1. Microfluidic device, strain and image acquisition

The microfluidic device used for the experiments shown here was developed by Mondragón-Palomino and colleagues [1]; soft lithography was used to make the devices as in [2]. *E. coli* MG1655 cells carrying the comparator construct described in [3] were loaded in the device as in [1]. For each experiment, time-lapse fluorescence microscopy experiments were performed using an inverted widefield fluorescence microscope (Leica) and images taken using an Andor iXon Ultra digital camera. The microfluidic device was enclosed inside an incubation chamber set to 37°C (Pecon). The microscope took every 5 minutes images of cells growing inside the microfluidic device trapping chambers. At each time point, a phase contrast image (PhC) and a green fluorescence image was acquired for each of the three trapping chambers. Green fluorescence images were used for the detection of GFP. Images were acquired using a 100 X objective. Exposure times were set to 100 ms for the PhC and green spectra.

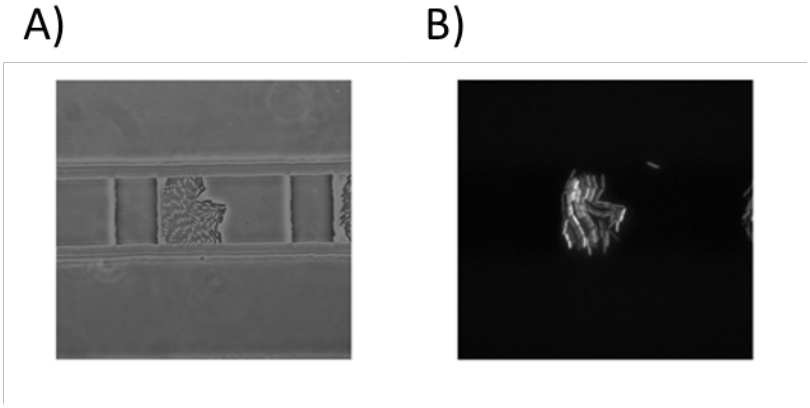


***Figure S1**. **Two channels frames**. A) Raw phase contrast image. B) Raw green fluorescence image.*

#### 2. Pre-processing of the raw images

For image pre-processing, firstly the user defines manually in the original image a crop; in so doing, the region of interest where cells are expected to grow is defined, excluding external objects (e.g. microfluidic device borders). Then, interpolation by *Interp2* is required to set the proper size of the image and to increase the resolution. Finally, filters such as *Wiener2* (2-D adaptive noise-removal filter) and *Medfilt2* (2-D median filter) are applied to reduce noise and blurriness (Fig. S2A, B); the *Adapthisteq* function is used to enhance the contrast of the grayscale image (Fig. S2C).

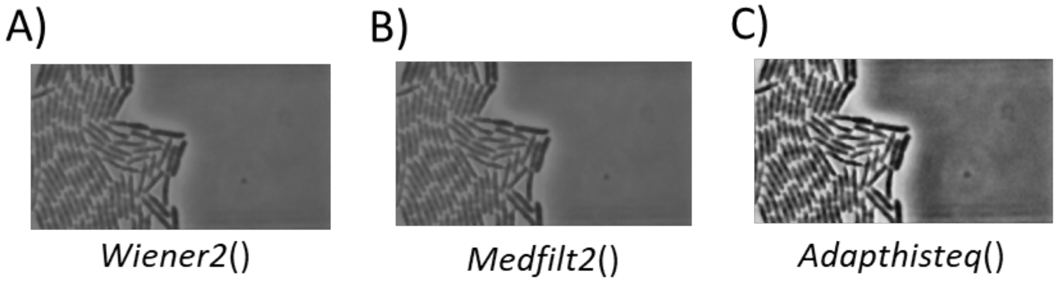


***Figure S2. Matlab filters for pre-processing.*** *A) Wiener2 filtering: adaptive noise-remove filter; B) Medfilt2 filtering: the function performs median filtering of a matrix in two dimensions; C) Adapthisteq contrast saturation: the function enhances the image contrast; it operates on small data regions (tiles), rather than on the entire image.*

### 3. Global and local thresholding

After pre-processing, the algorithm applies the Otsu method [4] to binarize the greyscale image by the *Imbinarize* function and uses morphological operators such *Imdilate* and *Imfill* to calculate the global area where cells are located (Fig. S3). Local thresholding is then applied to find cell centres and edges to segment individual cells by the *Niblack* and *Averagefilter* user-defined functions (Fig. S4).

#### a. Global thresholding

A first step for the computation of a mask is given by the identification of the "global area" where cells can be distinguished from the background. This rough result is achieved using three simple functions: *Imbinarize*, that calls the Otsu method and generates a binary image; *Imdilate* and *Imfill*, which dilate the objects in the image and then fill the internal gaps, respectively (Fig. S3).

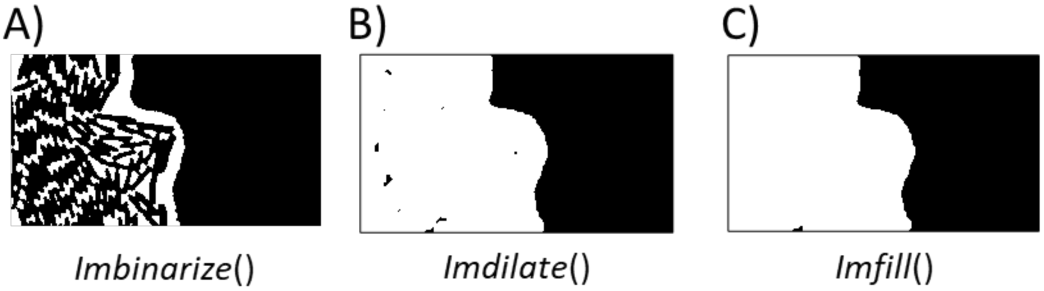


***Figure S3**. **Global thresholding functions**. A) The Imbinarize function binarizes the grayscale image by thresholding (Otsu method). The threshold is automatically defined as global, unless a different method is specified. B) The Imdilate function dilates the image referring to the shape of a structuring element object (STREL). C) The Imfill function performs a filling operation on background pixels of the input binary image specified.*

#### b. Local Thresholding

Local thresholding is needed to differentiate individual cells in a binary mask. First, an automatic crop is applied to select the zone of the image containing cells (Fig. S4A). The size for the crop is chosen referring to the mask size, previously calculated by global thresholding. To apply a local thresholding routine, we use two user-defined functions (*Niblack* and *Averagefilter*) to segment individual objects; the *Averagefilter* function is nested inside the *Niblack* call (Fig. S4B).

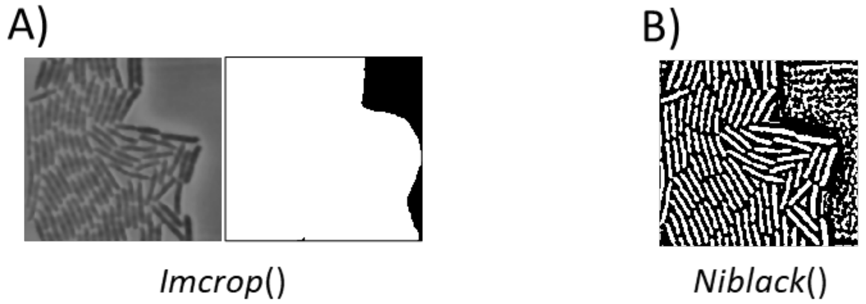


***Figure S4*. *Local thresholding functions.*** *A) The Imcrop function crops an image; if no area is specified, the command can create an interactive window to select an area to be cropped. B) Niblack and Averagefilter user-defined functions: these nested functions, based on the Niblack algorithm in* [5]*, divide foreground and background using local information about the image.*

### 4. Segmentation refinement process

After local thresholding, we refine the segmentation results to get rid of possible artefacts of the segmentation process. Firstly, small objects which are out of the individual cells size range are removed. This is done by a selective Matlab function – *Bwareafilt* - that keeps the *n* largest objects according to user-defined criteria (Fig. S5A). Then, the morphological operators *Imfill* and *Imopen* are called to smooth the edges and fix small objects which are likely to be artefacts of the segmentation algorithm (Fig. S5B and C). The resulting local thresholding mask is shown (Fig. S5C); there are still few small objects which are removed later by a user-defined function based on fluorescence calculation (see below).

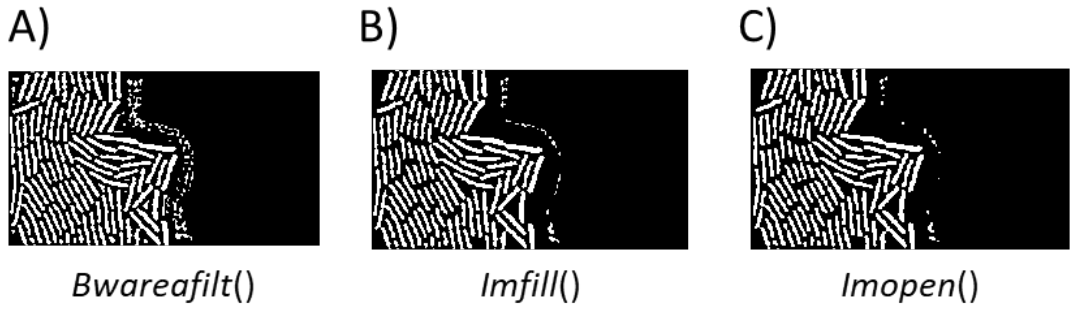


***Figure S5*. *Segmentation refinement functions and mask resulting from local thresholding*.** *A) The Bwareafilt function extracts objects from a binary image by size. A value for the size must be specified. B) Imfill function, as in Fig. S3C. C) Imopen function: the morphological*

*operation is an erosion followed by a dilation, using the same structuring element (STREL) for both operations. This operation allows a further removal of some small objects.*

A further refinement routine is carried out by the user-defined function *segmentationRefinement*. This method aims at removing artefacts due to the fact that individual cells, if too close, can get segmented as a single object. Firstly, the algorithm chooses the objects which have an area twice bigger than the average area calculated over all the objects present in the mask. These larger objects are further segmented until identifying individual cells (Fig. S6).

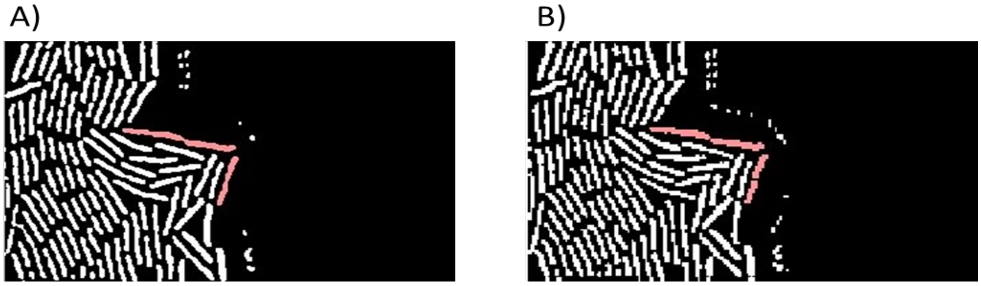


***Figure S6*. *Long objects refinement*** *A) The image shows the original mask before the refinement process. Long objects, highlighted in red, are detected by the algorithm. B) The bigger objects are divided by a line, drawn calculating the convex points representing edges where two cells touch.*

A final refinement of the mask is made by deleting the remaining small objects. The average cell area evaluated in the previous step is used again to identify objects with an area at least 50% smaller than the average. Through the user-defined function *SmallObjectsRemoval*, those objects are deleted from the mask, which is then resized to the original crop size defined in the pre-processing stage (Fig. S7).

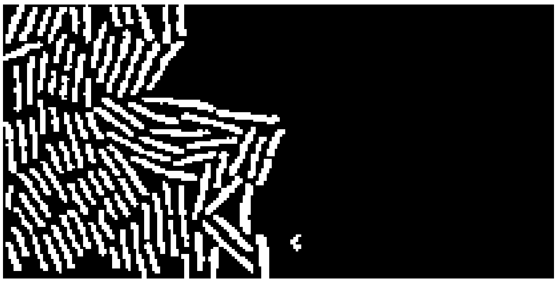

***Figure S7*. *Small objects removal*.** *The image shows the last process of small objects removal (user-defined function) to delete the spots around the segmented cells. This is also the final mask used to calculate the fluorescence.*

#### 5. Fluorescence calculation

The final mask (Fig. S7) is overlapped to the green field image to calculate the fluorescence using the custom function *fluorescence_eval.m*. This is computed as the sum of all pixels in the segmented area minus the background fluorescence value. The background is calculated as the mean of the pixels acquired cropping an area without cells from the green field channel.

A threshold fluorescence value (using *fluorescence_eval_Init.m*) is computed to exclude cells over a fluorescence threshold set by the user: in this way possible outliers (e.g. mutants resulting from phototoxicity effects) are removed. The threshold is calculated as the highest fluorescence in a set of images. The final average fluorescence value is then computed as the mean of the fluorescence exhibited by all the objects in the final mask after removing those over the threshold.

The algorithm outputs at each sampling time of the experiment the average fluorescence across the cell population (Fig. 1C, Supporting Movie 1), its statistical moments and the fluorescence histogram. The algorithm outputs also the cell number by counting the numbers of individual objects [6].

### Mammalian cell segmentation

The segmentation algorithm for mammalian cells, similarly to the bacteria one, can be applied to different types of images to distinguish the foreground from the background. The algorithm can identify cellular marks, compute fluorescence and track cells over time. The steps and parameters have been optimised for our experimental settings (see below); the user might need to refine parameters or could possibly skip some pre-processing steps depending on the acquired raw images and the cells used. The Matlab files implementing the reported steps are '*Main_Offline.m*', '*Offline_function.m*', '*Mask_n_Track.m*' and '*fluorescence_evaluation.m*'.

#### 1. Microfluidic device, cell lines and image acquisition

Experiments analysed in this paper were performed with either of two cells lines: a mouse embryonic stem cell (mESC) line stably carrying a destabilised GFP replacing the entire Rex1 coding sequence in one allele (Rex1-GFPd2) [7], or dual-reporter mESCs carrying coding regions for eGFP and mCherry downstream of transcriptional start sites for mir-302 and mir-290 clusters, respectively [8]. Cells were cultured on gelatinised tissue culture dishes at 37˚C in a 5% CO2 humidified incubator in Dulbecco's modified Eagles medium (DMEM; Sigma; D5796) supplemented with 15% fetal bovine serum (Sigma; F7524), non-essential amino acids (Gibco; 11140035), L-glutamine (Gibco; 25030024), sodium pyruvate (Gibco; 11360039), Penicillin-Streptomycin (Sigma; P4458), 2-mercaptoethanol (Gibco; 31350010) and 10 ng/ml mLIF (Peprotech; 250-02). Both cell lines were transfected with a plasmid to express a transgene to encode nuclear Histone 2B (H2B) tagged with infrared fluorescent protein (iRFP), as in [9].

The microfluidic device for mammalian cells was designed and characterised by Kolnik *et al.* [10]. A master mould of the device (MicruX technologies) was used to produce device replicas by PDMS moulding. Microfluidic device fabrication, processing and cell loading into devices were performed as previously described [9]. Cells were vacuum-loaded into microfluidic chips and pre-cultured with constant perfusion of fresh media while allowed to attach overnight in a humidified incubator; Rex1-GFPd2 cells were supplemented with 10µM cell permeable CellTracker blue (Invitrogen; C12881), and cultured in gelatine-treated chips; the microfluidic devices were precoated with fibronectin for dual-reporter mESC experiments. The device was secured onto a motorised microscope stage within an incubation chamber (incubator i8; Leica microsystems) and maintained at 37°C in a 5% $CO_2$. Time-lapse imaging was performed to acquire phase and fluorescent images using a Leica LASX live cell imaging workstation on a

DMi8 inverted fluorescent microscope with an Andor iXON 897 Ultra digital camera and a 40x objective (PlanFluor DLL 40x Ph2 Nikon). Adaptive focus control was employed on chosen fields and images were acquired every 60 mins. The set-up for the experiment in Fig. S8-S10 (i.e using the Rex1-GFPd2 mESC reporter line) consisted of 4-channels (Phase contrast, Green, Blue and Infrared) whilst the set-up for experiment in Fig. S11-S13 (i.e. using the dual reporter mESC line) consisted of 3 channels (Phase contrast, Red and Infrared).

2. **Selection of a channel for the segmentation process**

The algorithm applied to the first exemplar experiment shown here (Fig. S8-S10, Supporting Movie 2) uses an Otsu method to identify masks and quantify the fluorescent reporter expression in a dome-shaped population of Rex1-GFPd2 mESCs. We applied the Otsu method to either the phase contrast images, or images collected using a Blue channel to capture a blue dye incorporated into cells (Fig S8A, D). Blue dye images were used for segmentation given the superior contrast in the scale of grey as compared to phase contrast images; furthermore, if based on dye images, the algorithm avoids including in the foreground non-fluorescent objects which are not cells (e.g. edges of the microfluidic device). In what follows, we show only the segmentation process in case dye images are used; still, the same routine could be used for phase contrast frames.

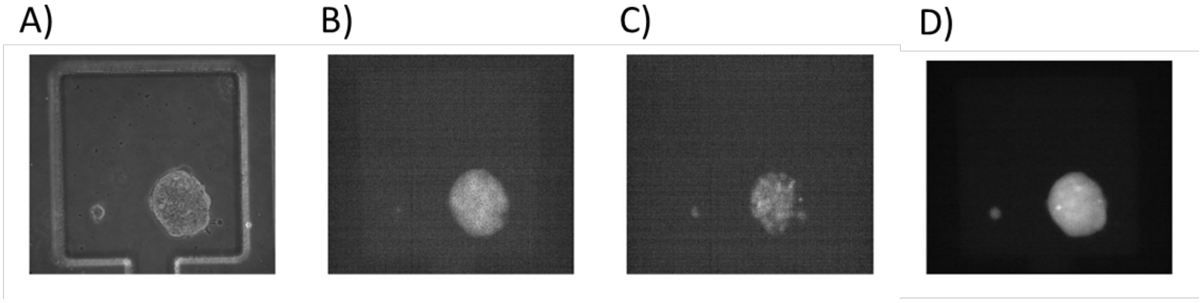


***Fig. S8*. *Raw images, Rex1-GFPd2 mESCs.*** *A) Phase contrast image. B) Green fluorescence image. C) H2B- Nuclei tag image. D) Blue dye image.*

3. **Pre-processing of the raw images**

As for bacterial cells, at the start of the time-lapse a crop is manually made to define the region of interest from the initial picture frame (Fig. S9A); the same crop is then used throughout the time-lapse. At the start of the experiment, an additional crop is required to identify a region without cells, which will be later used to calculate the background fluorescence.

Then, the cropped image with cells is filtered to adjust the colour intensity values; 1% of data is saturated at low and high intensities by the *Imadjust* MATLAB function (Fig. S9B).

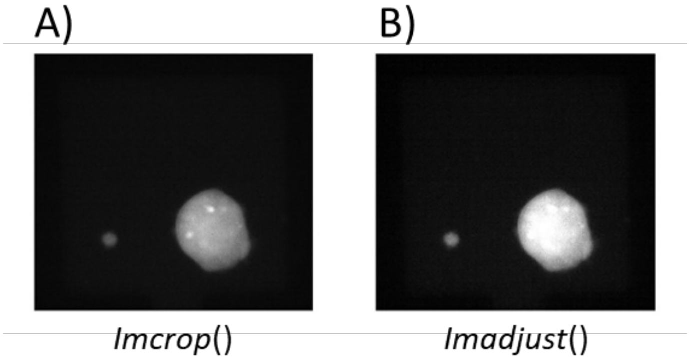


*Fig. S9. Pre-processing functions. A) Imcrop function: the original raw image is manually cropped from the first timeframe. The crop function is the same described in Fig. S4A. B) Imadjust function operates a saturation of 1% of data at low and high intensities.*

### 4. Global thresholding and single-cell tracking

#### a. Global Thresholding

Before and after applying the Otsu algorithm (*Imbinarize* function), some filtering actions might be required to improve the image and mask quality, as for bacteria cells (Fig. S10). Of note, parameters in these functions (e.g. the structuring element object within the *Imerode* function) need to be optimised by trial and error, depending of the images; if segmenting images acquired under different settings, some of these functions might be redundant.

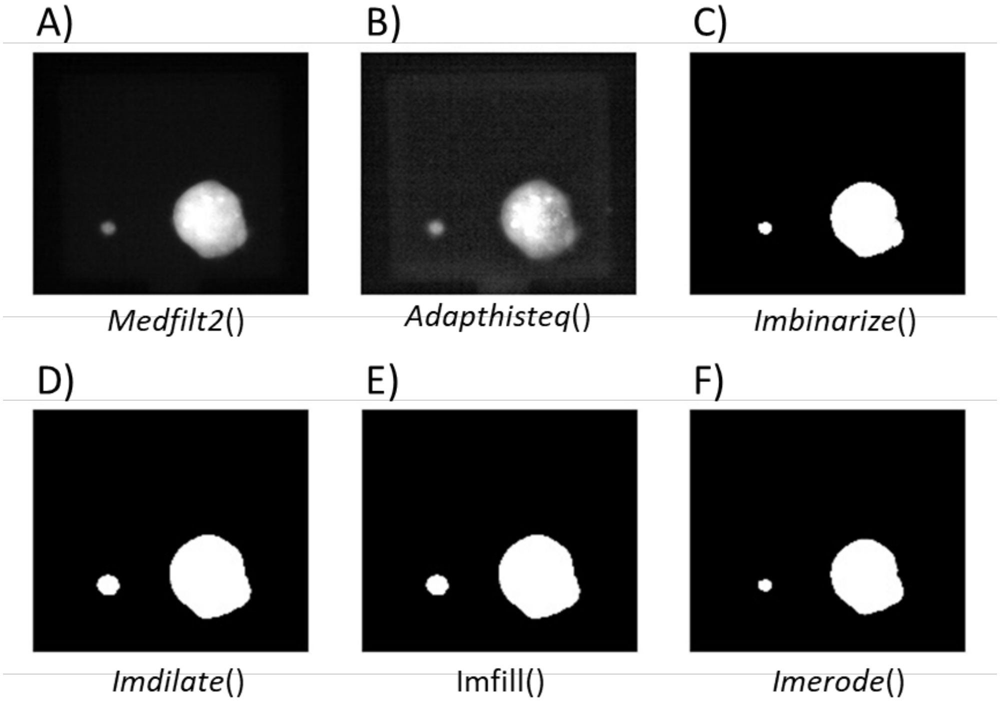


*Fig. S10. Global thresholding and filters. A) Medfilt2* filter: as described in Fig. S2B, this function performs a median filtering of a matrix. B) Adapthisteq* function: as described in Fig. S2C, enhances the contrast of images. C) Imbinarize function: as described in Fig. S3A. D) Imdilate* function: as described in Fig. S3B. E) Imfill* function: as described in Fig.*

S3C. F) Imerode* function: erodes images referring to the shape of a structuring element object (STREL). * Depending on the raw images, some of these functions might be redundant.

#### b. Single-cell tracking

Single-cell tracking is another feature embedded in our code, always relying on the Otsu method for mask recognition. Here we show an application to a time-lapse where dual-reporter mESC images of three channels were collected (Fig. S11); to clearly identify individual cells we used an H2B-tag reporter for nuclei (Fig. S11C).

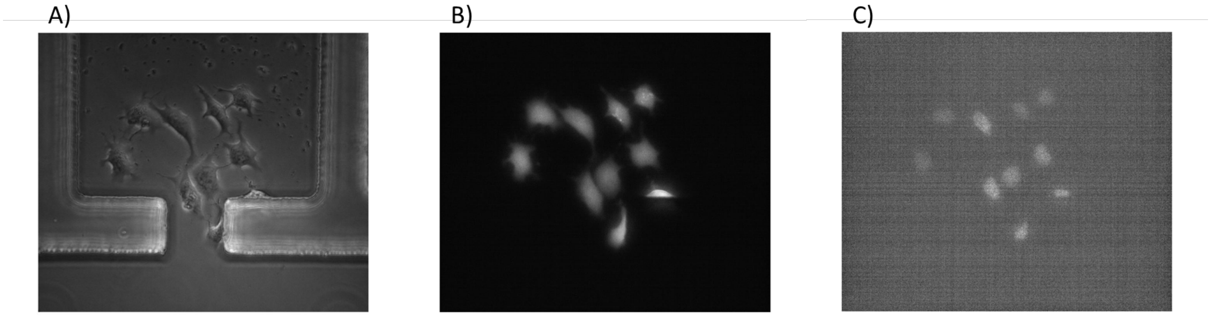


***Fig. S11. Raw images of dual-reporter mESCs used in single-cell tracking experiment.*** *A) Phase contrast image. B) mCherry fluorescence image. C) H2B nuclear tag image.*

The tracking code firstly computes a rough mask detecting nuclei intensity by applying filters and thresholding functions (Fig. S12), as defined within the user-defined function *mask_n_Track*. Then, the mask is used to identify and label connected objects (i.e. individual cells). We employed the mCherry fluorescent image (expressed both in the nucleus and in the cytosol, Fig. S11B) to calculate the average cell area; using the *Bwareafilt* function, objects whose area is less than the 5% of the average are removed (Fig. S12F). This step might be neglected if the nuclei images have a higher contract as compared to the background than in our exemplar experiment.

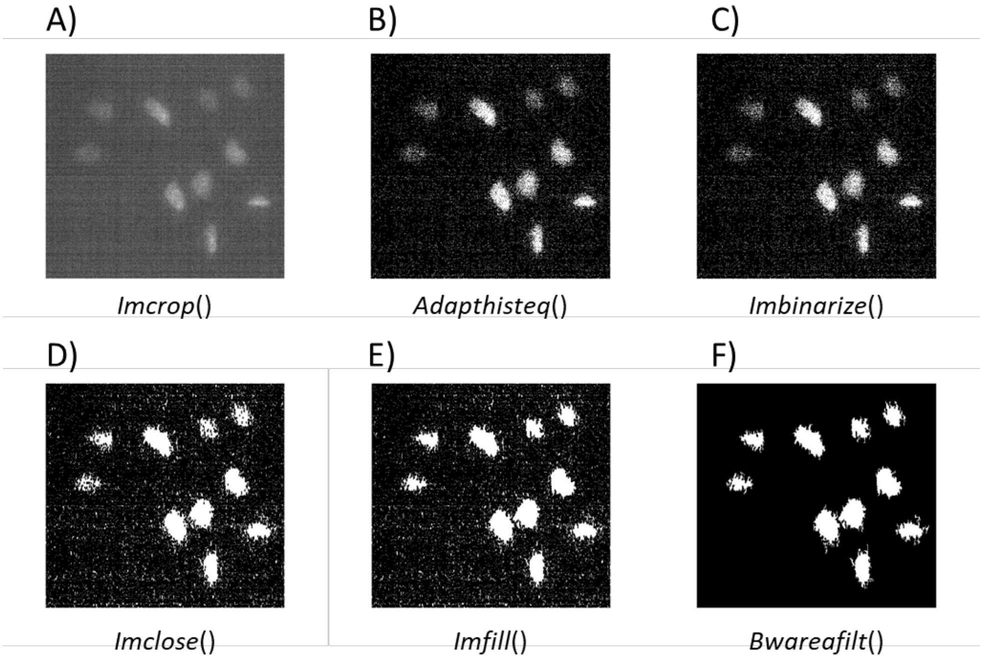

*Fig. S12. Filtering functions. A) Imcrop function: the original raw image is manually cropped in the first timeframe. The crop function is the same described in Fig. S4A. B) Adapthisteq function: as described in Fig. S2C. C) Imbinarize function: as described in Fig. S3A. D) Imclose function: the morphological open operation is a dilation followed by an erosion, using the same structuring element (STREL) for both operations. E) Imfill* function, as described in Fig. S3C. F) Bwareafilt function: as described in Fig. S5A.*

Using the Matlab function *Bwlabel*, the code then associates to each object a label, that is going to be maintained throughout the whole time-lapse experiment. For each object, a centroid is evaluated using the Matlab function *Regionprops*; this function also computes the major and minor axis of the ellipses defining the dimensions, their orientation and the area of cells nuclei. The algorithm then performs tracking by using the custom functions *MaskGenerator* and *Tracking_fun* (the latter encompassing *SaveStructEllipse, TraceOldNew* and *UpdateOBJS* functions).
*MaskGenerator* generates multiple masks, one for each cell detected, overlapping the rough mask previously computed with ellipses data generated by *Regionprops*.
Then, from the second timeframe onwards, the centroids among contiguous timeframes are compared. Each object (i.e. cell) is saved by *SaveStructEllipse* along with the ellipse associated data and computed masks, and a minimum distance problem is solved by the *TraceOldNew* function to identify an object in two consecutive frames. The label associated to each cell is kept or swapped with another cell during the experiment according to the *updateOBJS* method; a cell label is removed only in case the cell goes out of the focus or leaves the cropped region. Some parameters of the routine (e.g. the maximum nuclei dimension) need to be fixed by the user given the acquisition settings and specific cell morphology. In Fig. S13 we show two consecutive frames and the relative tracking of individual cells.

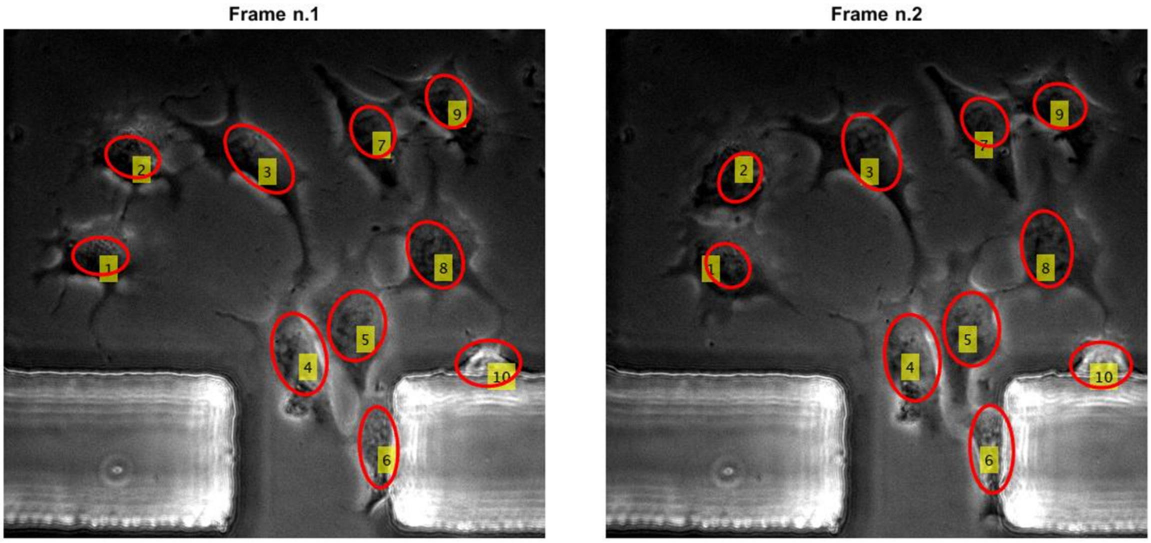


***Fig. S13. Cell tracking.** The tracking of individual cells using the functions Mask_n_Track and Tracking_fun in two consecutive frames is shown.*

### 6. Fluorescence calculation

Once a mask is available, it can be applied to the frame of interest to evaluate the associated fluorescence value at cell population (Fig. 1D, Supporting Movie 2) or single-cell (Fig. 1E,

Supporting Movie 3) level; an average background fluorescence is subtracted to cell fluorescence. The background intensity value can be either computed by cropping an empty area of the frame or by defining the reversed logical mask. In the latter case, the background is then obtained as it would be for cells fluorescence, i.e. by applying the mask to a frame and averaging over the non-covered area. When a fluorescent tag is inserted into mammalian cell nuclei, a nuclei normalization can also be performed within fluorescence computation.
